# Placing ancient DNA sequences into reference phylogenies

**DOI:** 10.1101/2020.12.19.423614

**Authors:** Rui Martiniano, Bianca De Sanctis, Pille Hallast, Richard Durbin

**Affiliations:** Department of Genetics, University of Cambridge, Cambridge CB2 3EJ, UK; School of Biological and Environmental Sciences, Liverpool John Moores University, Liverpool L3 3AF, UK; Department of Zoology, University of Cambridge, Cambridge CB2 3EJ, UK; Institute of Biomedicine and Translational Medicine, University of Tartu, 50411 Tartu, Estonia; Wellcome Sanger Institute, Cambridge CB10 1SA, UK

## Abstract

During the last decade, large volumes of ancient DNA (aDNA) data have been generated as part of whole-genome shotgun and target capture sequencing studies. This includes sequences from non-recombining loci such as the mitochondrial or Y chromosomes. However, given the highly degraded nature of aDNA data, post-mortem deamination and often low genomic coverage, combining ancient and modern samples for phylogenetic analyses remains difficult. Without care, these factors can lead to incorrect placement.

For the Y chromosomes, current standard methods focus on curated markers, but these contain only a subset of the total variation. Examining all polymorphic markers is particularly important for low coverage aDNA data because it substantially increases the number of overlapping sites between present-day and ancient individuals which may lead to higher resolution phylogenetic placement. We provide an automated workflow for jointly analysing ancient and present-day sequence data in a phylogenetic context. For each ancient sample, we effectively evaluate the number of ancestral and derived alleles present on each branch and use this information to place ancient lineages to their most likely position in the phylogeny. We provide both a parsimony approach and a highly optimised likelihood-based approach that assigns a posterior probability to each branch.

To illustrate the application of this method, we have compiled and make available the largest public Y-chromosomal dataset to date (2,014 samples) which we used as a reference for phylogenetic placement. We process publicly available African ancient DNA Y-chromosome sequences and examine how patterns of Y-chromosomal diversity change across time and the relationship between ancient and present-day lineages. The same software can be used to place samples with large amounts of missing data into other large non-recombining phylogenies such as the mitochondrial tree.

## Introduction

The development of high-throughput sequencing methods and their application to archaeological remains has dramatically changed our understanding of deep human history. Alongside approaches using autosomal loci, the study of Y chromosomes and mitochondria has provided valuable insights, both because of the resolution of the phylogeny and also because they provide information about sex-biased migrations, kinship and social systems [1, 2, 3]. This is due to the fact that they are transmitted from a single parent, so avoiding recombination and leading to a simple tree phylogeny. However, there are substantial challenges associated with the analysis of aDNA in a phylogenetic context, especially for the Y chromosome because of its larger size (approximately 10Mb of callable sequence) [4] comparatively to the mitochondria (∼16 kb), as well as its lower copy number number so data are typically lower coverage.

The highly degraded nature of aDNA data, including short fragment size, post-mortem deamination and high fractions of missing genotypes [5, 6, 7], can lead to errors in variant calling and to incorrect placement of ancient DNA sequences within a phylogeny [8]. In particular, many standard phylogenetic methods require significant overlap of genotypes across samples, which is unfeasible when analysing a large number of samples simultaneously [9].

Another important factor which complicates phylogenetic analysis of ancient Y-chromosomes is that it derives from two distinct sources: one is whole-genome shotgun (WGS) sequencing, which provides data across the whole Y-chromosome sequence, and the other consisting of targeted approaches, where biotinylated probes are designed to be complementary to specific genomic regions or markers, and are subsequently retrieved through the use of magnetic beads and sequenced. A common example of the latter is the 1240k capture array [10], which targets known polymorphic sites along the genome. Both of these approaches have their advantages and disadvantages: SNP-based targeted approaches, despite suffering from ascertainment bias, are cost-effective especially when applied to less well-preserved remains with a lower fraction of endogenous DNA; WGS or region-based targeted capture, on the other hand, are relatively unbiased and allow obtaining novel variation, but may be prohibitively expensive to obtain high coverage data and therefore are often low coverage with a large fraction of sites without any genotype information [11]. While there are methods which use likelihoods for the placement of sequences into a pre-estimated phylogenetic tree, such as pplacer [12] and RAxML’s Evolutionary Placement algorithm [13], these do not take the degraded nature of ancient DNA into account and can provide erroneous assignments. They are also computationally expensive when applied to the thousands of samples available now for analysis [14, 15].

Sequencing of the non-recombining portion of the Y-chromosome has enabled the rapid and unbiased discovery of new Y-chromosome variants. The International Society of Genetic Genealogy (ISOGG; https://isogg.org/tree/) has been cataloguing new informative Y-chromosome variants during the last 15 years, and currently lists approximately 73,000 unique biallelic variants with different levels of confidence. However, curation of new variants is time-consuming and problematic: it can take years until variation from new sequencing studies is added to the ISOGG database and despite major effort a considerable subset of variants only have provisional assignments to specific Y-chromosome lineages, or even contain errors which then need revision.

With these aspects in mind, studies such as [16] and [17], have examined allele status in ancient samples at specific branches of large modern Y-chromosome trees such as from the 1000 Genomes project. By including both novel and known mutations, these studies increased the probability of a given ancient sample having reads overlapping informative branch-defining positions, as noted by [14]. However, no publicly available automated way of doing this exists.

Here we provide software and an associated workflow, pathPhynder, for integrating ancient DNA data from multiple sources (target capture and shotgun sequencing) and of variable genomic coverage into present-day phylogenies. To increase its specificity, pathPhynder supports updating and expanding the reference tree and panel of known variants by adding present-day Y-chromosome or mtDNA sequences, for example from newly sequenced diverse populations, to maximize the probability of overlap with sparse aDNA sequences and increase lineage informativeness. Furthermore, pathPhynder also provides a visualization tool which allows inspection of the number of markers in support of or conflict with aDNA sample assignment to each branch. While handling errors and missing data correctly by working in a likelihood framework, pathPhynder is also computationally efficient, scaling linearly with both sites and samples and taking only a fraction of a second to place a query into a large tree.

We apply pathPhynder to place previously published African aDNA samples of variable coverage into a present-day Y-chromosome tree. This increases the resolution beyond that in the original publications, revealing more detailed relationships between prehistoric and modern lineages and how they have changed through time.

## Results

### Description of the Method

The inputs to pathPhynder are a pre-existing reference phylogeny in standard newick format, a reference VCF file containing the genotypes of the individuals in the phylogeny, and BAM files of the ancient DNA query samples mapped against the same reference genome sequence as the phylogeny VCF (or alternatively an already processed VCF file of the query samples).

The pathPhynder workflow is represented in Figure 1. The first step assigns informative SNPs from the reference VCF to each branch of a phylogeny estimated with high-coverage individuals. This can be achieved by using the ‘phynder’ software, which estimates the likelihood of each biallelic SNP at each branch of the tree. These variants and their location at tree branches can then serve as an initial guide for placing aDNA samples, and for visualisation.

**Figure 1:**
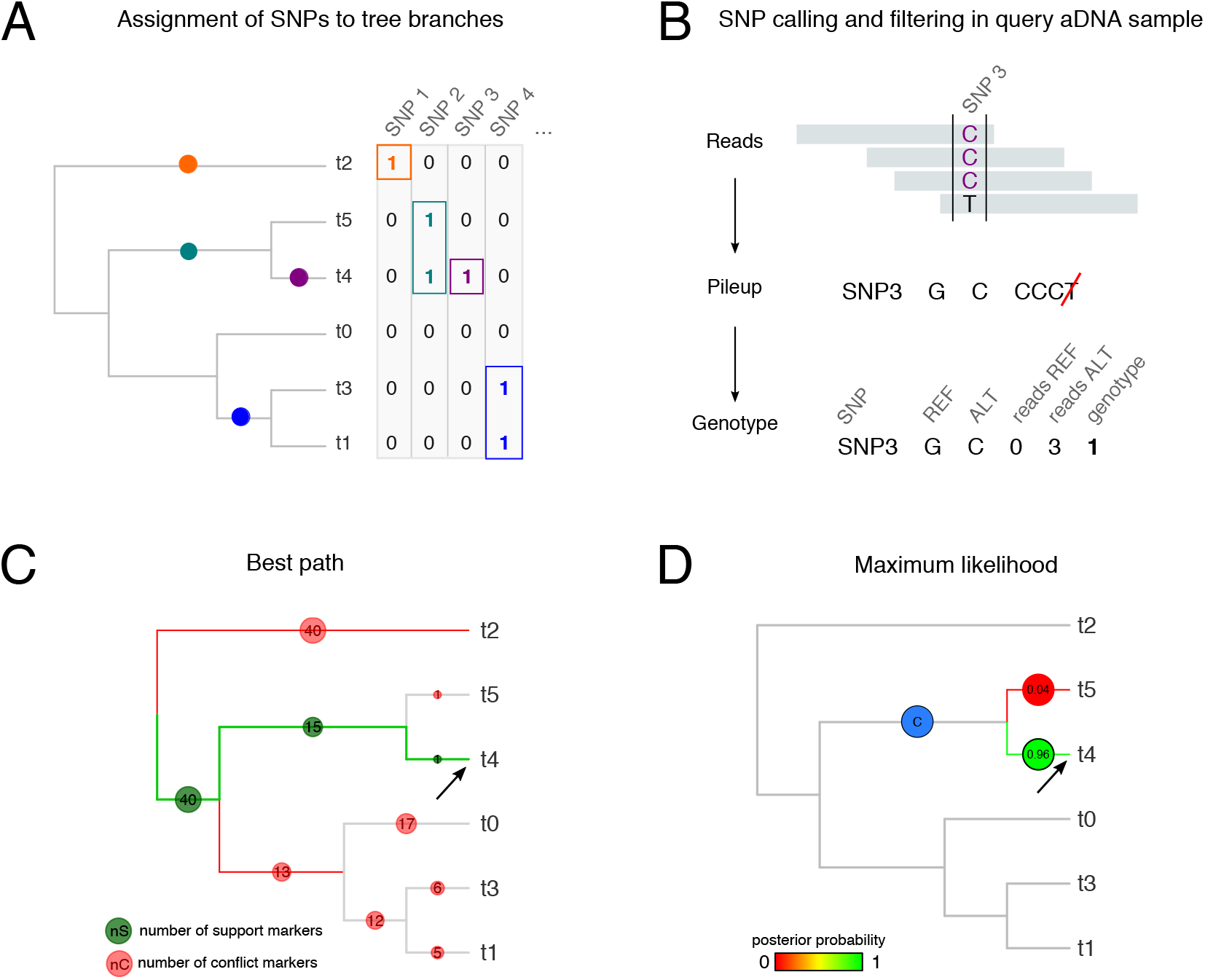
Overview of pathPhynder workflow. We illustrate the method using a small simulated dataset of 6 reference samples (tips t0 to t5) and 112 SNPs. (A) The initial step is assignment of phylogenetically informative SNPs in the reference dataset to branches. This can be achieved with phynder by estimating the likelihood of each SNP at any given branch of the tree. (B) A pileup from aDNA reads of the query aDNA sample is generated at each SNP, then filtered for mismatches and potential deamination. Here, SNP3 is covered by four reads, three ‘C’s and one ‘T’. Because SNP3 is defined by alleles G and C, the T base is excluded as likely to be caused by post-mortem deamination. (C) Best path method: aDNA sample genotypes for each SNP are assigned to the corresponding branch of the tree and binned into support and conflict categories. All possible paths from root to tips are traversed, evaluating the number of markers in support (green) or conflict (red). The best path (green edges), is the one containing the highest number of support markers (in this case, 56 markers). (D) Maximum likelihood method: the likelihood of the entire tree is calculated by placing the query aDNA sample on each edge of the phylogenetic tree. The likelihoods are converted to posterior probabilities using Bayes’ rule and the sample is placed on the branch with the highest posterior, with other branches with posterior probability greater than 0.01 also indicated. The likelihood method also finds the lowest branch in the tree for which the sum of posterior probabilities for the whole clade below that branch (including the branch in question) is greater than 0.99 (shown as a blue circle with a ‘C’), 15 which provides a conservative assignment when placement is uncertain. The arrows point to the correct location for the phylogenetic placement of the query aDNA sample.

Next, a pileup of base calls at the informative sites identified in the previous step is generated for each ancient sample using samtools (by default requiring base quality at least 20 and mapping quality at least 25), and subsequently filtered for error and deamination as follows (Figure 1B). First, base calls matching neither of the REF or ALT allele are removed. Then three further filtering modes are available: ‘no-filter’, where all remaining calls are retained; ‘default’, in which singleton T calls at C/T sites and singleton A calls at G/A sites are removed to account for possible deamination, and finally, ‘transversions’, which excludes all transition (C/T and G/A) SNP sites from analysis. Following this the genotype is called as the most frequent base so long as it is present at least a set fraction (default 70%) of the remaining base calls; otherwise the genotype is set to missing. An alternative option is to call genotypes at known informative SNP sites using external software and then to pass them to the program as a VCF file.

For the sequence placement step, the user can choose between two distinct methods: best path or maximum likelihood. In the best path method (Figure 1C), the SNP counts for a given aDNA sample are assigned to the respective branches, and we traverse possible paths from root to tip in the tree systematically. During this process, if a branch contains a number of conflicting markers greater than a user defined maximum threshold (default 3), the path is stopped and the next one is considered. The path containing the highest number of supporting markers is chosen as the best. This method is akin to the one implemented in yhaplo [18] but applies to all SNPs in the reference samples, rather than the ISOGG pre-curated SNP set. The number of mismatches observed at the assigned branch for a given sample is used to estimate where along the branch the sample should be inserted.

In the likelihood method (Figure 1D), we first score the likelihood of placing the query sample on each branch of the tree under a simplifying assumption that ignores mutations on that branch, then searches within the optimal branch using a full likelihood model that allows for repeated mutation at a site across the tree and data errors. The approximation is conservative. In addition to identifying the most likely branch, this approach provides Bayesian posterior probabilities for branches with posterior above a user defined threshold *p* (default 0.01) and the lowest branch for which almost all (1*− p*) of the posterior probability lies on or below the branch. Further details are provided in Supplementary Text 1.

### Method performance

We prepared a Y-chromosome dataset of 2,014 individuals from genetically diverse populations with genotypes at a total of 121,966 SNPs (Figure 2A, Material and Methods). We built a phylogeny from these samples using RAxML. Phynder assigned 120,908 SNPs (99.65% of the total) in the reference VCF file to the branches of the reference phylogenetic tree. A small number of SNPs (n=1058) were dropped, either because they were multiallelic (n=631) or because they could not be assigned with confidence (n=427), more precisely because their log likelihood was below a threshold, due to repeated mutation, repeated genotype error, or possibly errors in the tree topology.

**Figure 2:**
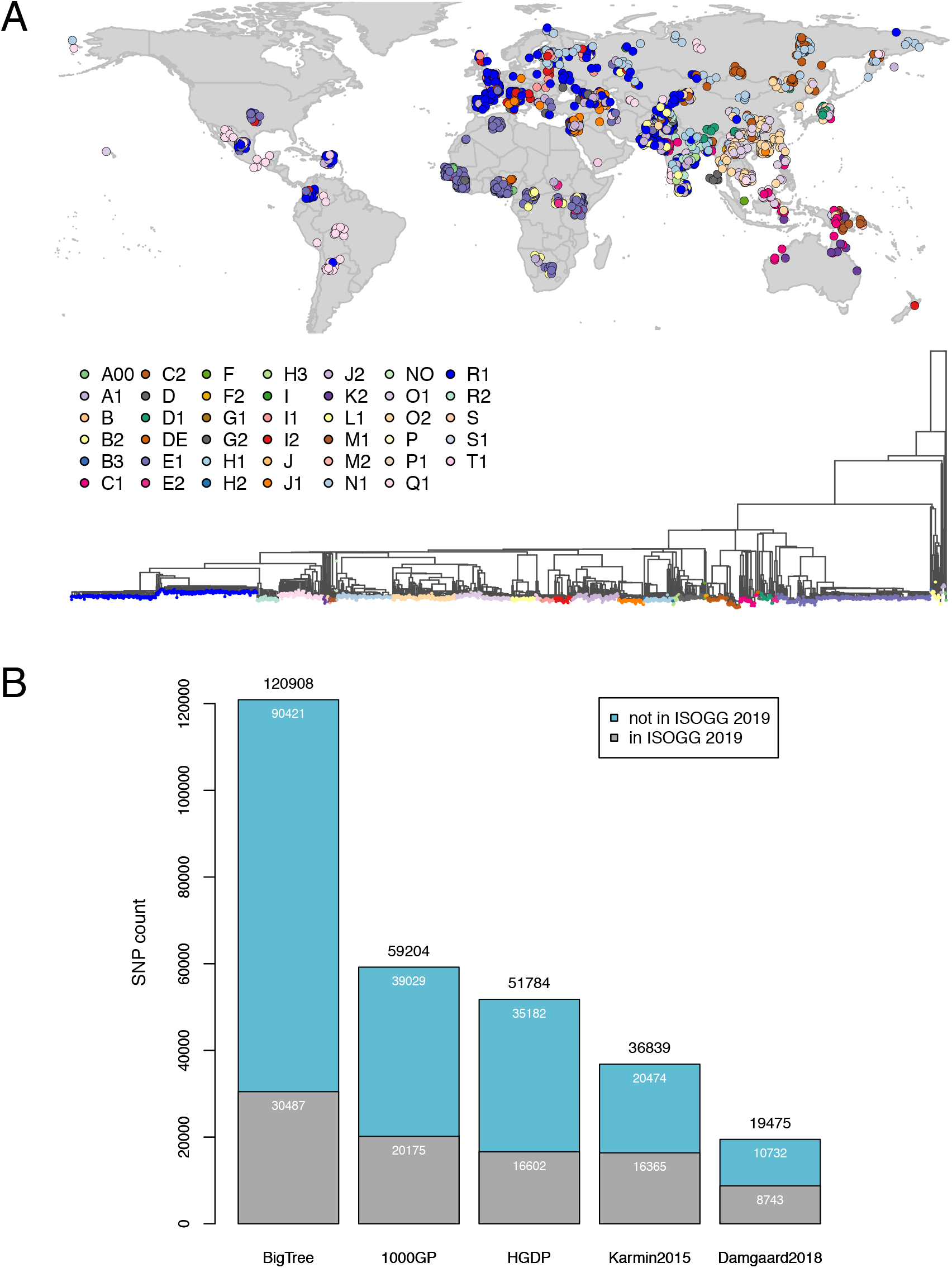
Overview of Y-chromosome SNP variation. A) Phylogenetic tree of the ‘BigTree’ Y-chromosome reference dataset which we compiled in the present-work and sample location (slightly jittered). B) Total SNP count of different datasets, distinguishing variants that have not yet been included in the ISOGG 2019-2020 database.

We compare the variation present in this dataset with previously published present-day data for the Y chromosome [14, 19, 15, 20, 21] by estimating the number of phylogenetically informative SNPs which can be assigned to branches of each tree (Fig 2B). Not only does this new data set contain more variants than all the others (n=121,335), but also the majority (n=90,421, 75%) have not yet been catalogued in ISOGG 2019-2020.

To evaluate the advantages of using this additional variation, we selected samples which had been assigned in the literature using catalogued variants in the ISOGG database to upstream branches of the phylogeny, such as BT-M91 or CT-M168, or were unassigned, and reanalyzed them with pathPhynder. In Figure 3 we show the distance between the previous and the newly assigned nodes. In most cases pathPhynder is able to use additional, uncatalogued variation in our new tree to improve the resolution of Y-chromosome lineage assignment (see also Supplementary Table S1).

**Figure 3:**
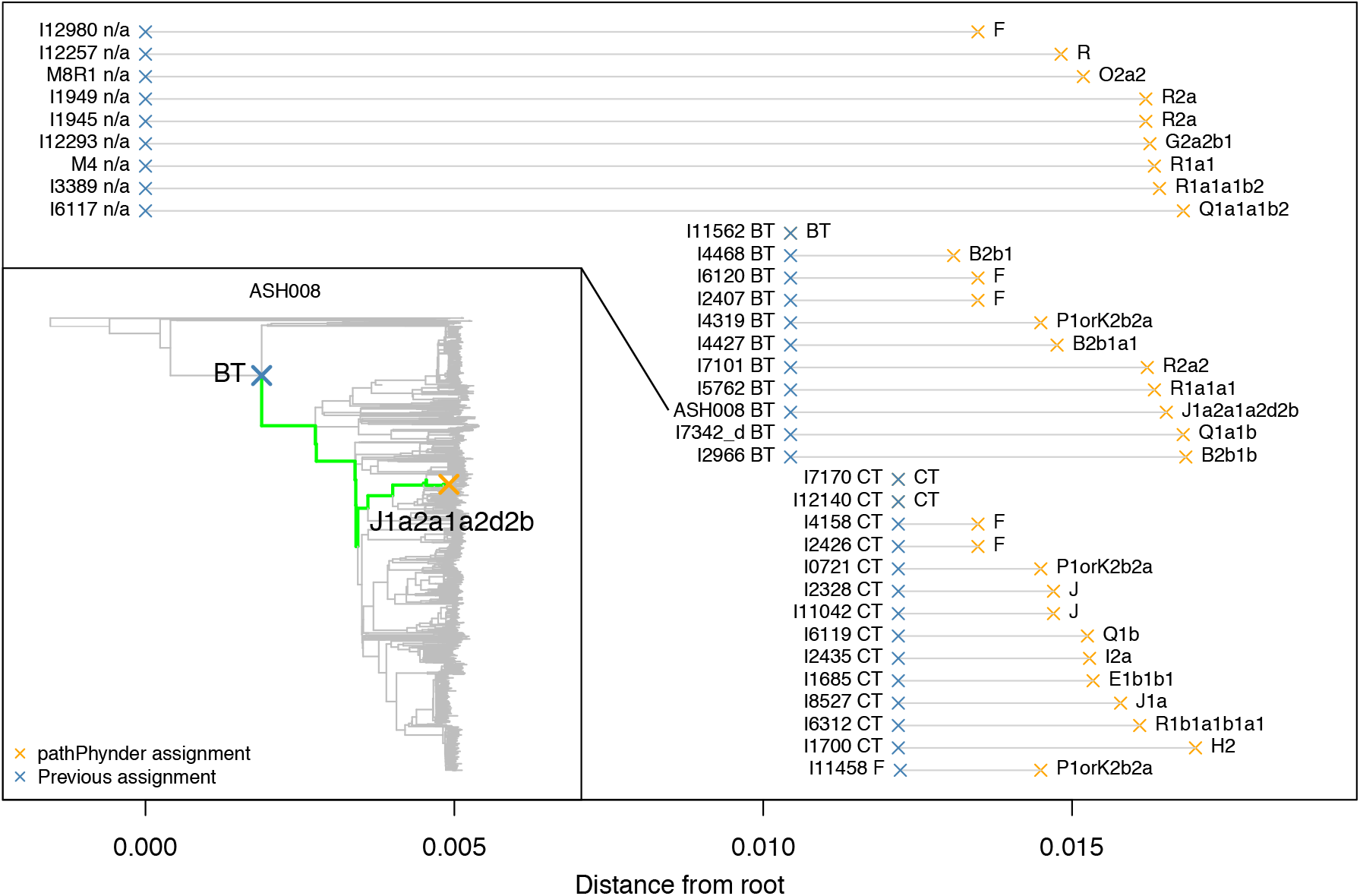
pathPhynder improves Y-chromosome lineage resolution when compared to standard haplogroup determination methods. We identified 34 low coverage samples which were assigned to higher level branches of the Y-chromosome tree in the literature (represented with blue crosses), and reassigned these with pathPhynder using the ‘BigTree’ dataset as a reference (orange crosses). We estimate the distance between the previous and the newly estimated nodes (grey lines connecting the crosses), showing that in most cases, pathPhynder can make use of additional, uncatalogued variation to improve the resolution of Y-chromosome lineage assignment. The phylogenetic tree (inset) provides an example of this process for sample ASH008, which was previously assigned to BT [49]. By making use of both catalogued and uncatalogued SNPs, this sample can now be placed in the J1a2a1a2d2b clade.

We next compared our approach to RAxML EPA (Evolutionary Placement Algorithm), an existing likelihood-based method for query sample placement in pre-estimated phylogenies. For this, we selected three individuals from our dataset belonging to the Biaka, Bedouin and Brahui populations who carry different Y-chromosome lineages (B2b1a1c2a2 ∼ -B26, J1a2a1a2d2b2b2c4b1c3a1a1 ∼-ZS5599, R1a1a1b2a1a1a1f∼ -Y928, respectively). We introduced missing genotypes at proportions ranging from 0.5 to 0.95 of the total data (∼120,000 SNPs), making a total of 30 query samples for placement in the phylogenetic tree. Prior to the query sample placement we excluded the three individuals from the reference tree. We then used pathPhynder’s best path, phynder’s maximum likelihood and RAxML’s EPA to place the query samples into the tree and compared the obtained result with the known true position in the original tree.

All three methods were similarly accurate and were able to place the query samples in the correct tree location between 77%-80% of the time (Supplementary Figure S1), with errors larger than a node distance of one only occurring beyond 85% missingness (∼ 18,000 SNPs) (Supplementary Figure S3). Interestingly, in the worst case of 95% (∼ 6,000 SNPs) missing data both likelihood methods (pathPhynder likelihood and RAxML) assigned to the wrong tips of the tree, perhaps due to these being longer and so having a larger prior in the absence of other data, whereas pathPhynder best path is more conservative, assigning to a higher node in the tree that includes the correct tip (Supplementary Figure S2). The ‘clade’ option of pathPhynder likelihood does essentially the same thing.

These results were unfiltered for deamination because they used present-day data. If we take the 80% missing point and add 10% deamination errors at transitions to simulate ancient DNA, then using the unfiltered data set the performance degrades substantially for likelihood approaches, with mean node error distances increasing to 27.67 (Supplementary Figure S4). With the ‘default’ filtering, which removes potential deamination, the error returns to 0. With the transversion-only filter it increases again to 3, because too many sites are dropped. The best path approach is much less affected by deamination because it stops traversing the path when it encounters more than 3 variants (default parameter that can be modified by the user) in conflict with query sample membership at a given branch. We note that straightforward application of standard tools such as RAxML will be equivalent to the no-filtering approach, and that a simplistic approach of restricting to only transversion sites leads to loss of valuable information.

In terms of speed, phynder’s maximum likelihood method is much faster than the other two, taking only 1 minute and 27 seconds for placing 30 query samples into our tree. pathPhynder’s best path method took 16 minutes and 56 seconds, and RAxML took approximately 76 minutes and 1 second.

### Ancient Y-chromosome diversity in Africa

In order to demonstrate the usefulness of our method for real data, we examine ancient and present-day Y-chromosome diversity in Africa by placing all ancient male samples from the African continent published at the time of this study (n=63) [17, 22, 23, 24, 25, 26, 27, 28, 29], and additional samples from the Levant (n=15) [30] into the Y-chromosome tree using pathPhynder.

As expected, the vast majority of the ancient African samples were placed into the A, B or E clades of the Y-chromosome tree (Figure 4 and Figure 5), substantially increasing the lineage resolution of 18 samples (Supplementary Table S2). The most ancestral human Y-chromosome lineage in our dataset is A00-L1284 [31], which is carried by two Mbo individuals from Western Cameroon [21]. Here, we establish that these two Mbo individuals in fact belong to the A00b-A4987 lineage. In the ancient DNA record, a single representative of A00 lineages has recently been identified in Shum Laka Cave, also located in the Cameroon, dating from ∼ 8 kya [26]. pathPhynder placed the Shum Laka sample on the edge leading to the two A00b individuals (Figure 4) with 2073 SNPs supporting this placement and 209 other in conflict (i.e. for which the Shum Laka sample carries the ancestral allele, including all five of the seven SNPs which ISOGG use to define the A00b lineage for which Shum Laka has data).

**Figure 4:**
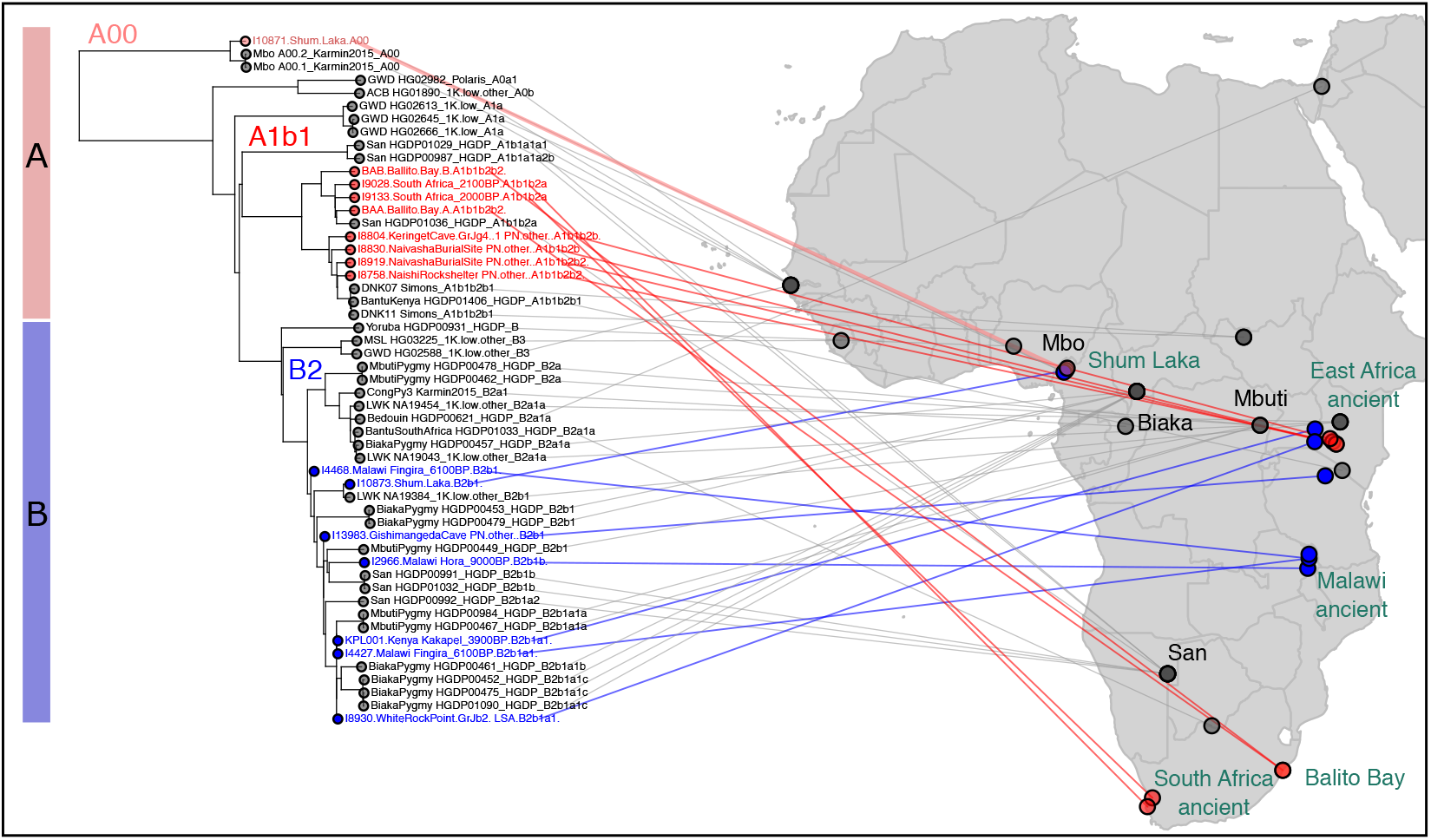
pathPhynder placement of ancient African samples into the A and B clades of the Y-chromosome tree. A and B lineages are mostly composed of present-day San, Mbuti and Biaka Pygmy populations. In terms of ancient DNA samples, these mostly belong to hunter-gatherer groups.

**Figure 5:**
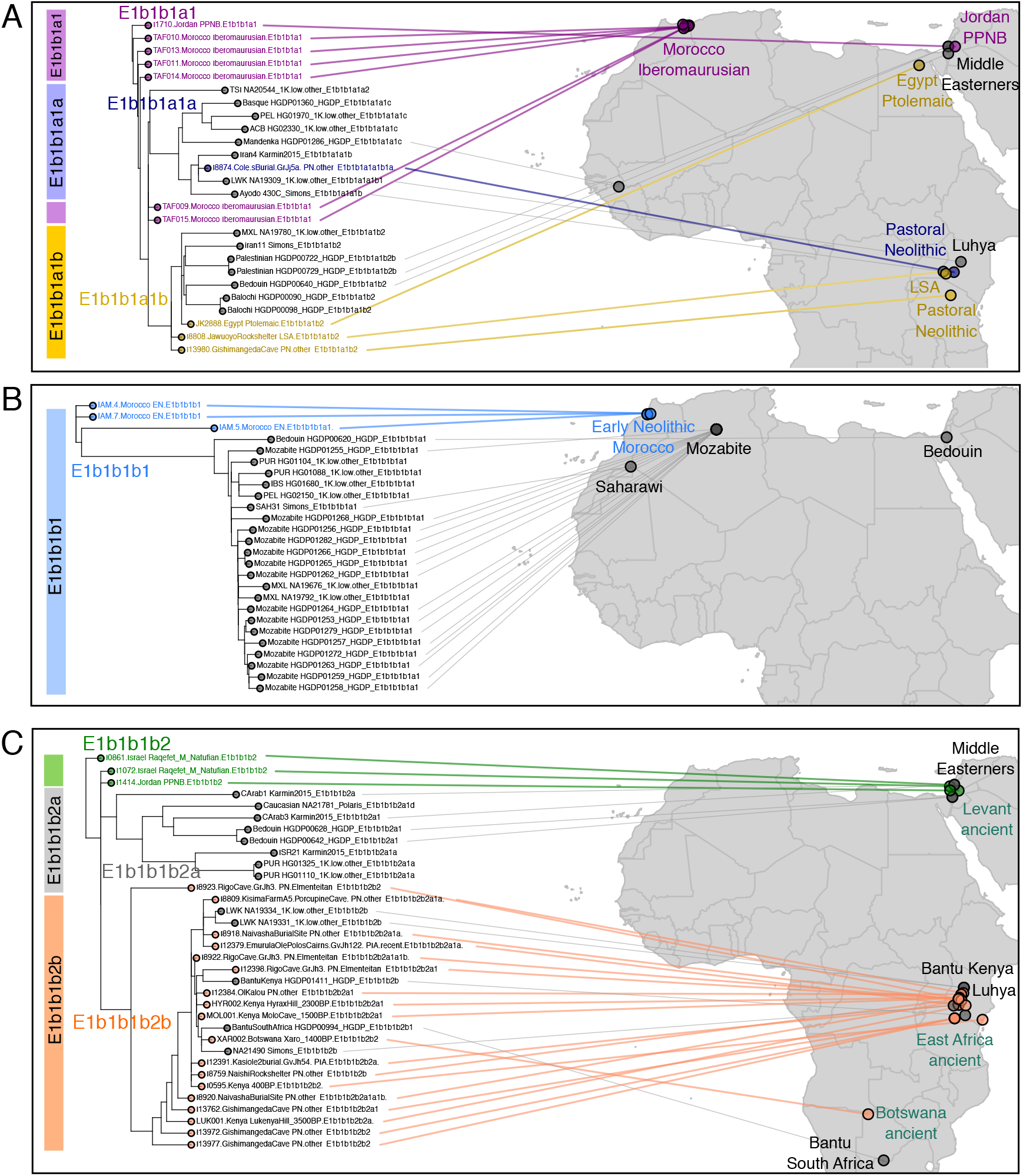
Placement of ancient African samples into the E subclades of the Y-chromosome tree. A) E1b1b1a1 lineages carried by Morocco Ibemaurusian period samples and one Jordan PPNB individual. B) E1b1b1b1 lineages mostly present in Algerian Mozabite populations and shared with Moroccan Early Neolithic samples. C) E1b1b1b2 lineages present in Pastoral Neolithic samples from East Africa and Levantine Natufians to whom they are ancestrally related.

Regarding clade A1-P305, it splits into A1a-M31, present in the Gambian Mandinka, and A1b-P108, present in the South African San and in the Dinka, a Nilotic group from Sudan. The pathPhynder placement revealed a strong geographical pattern: four approximately 2000 year-old South Africans from Balito Bay [27] and the Western Cape [22] were positioned in the A1b1b2a-M51 clade together with with a South African San individual, while East African Pastoral Neolithic (PN) samples from Kenya (Keringet Cave and Naivasha burial site) were placed in the A1b1b2b-M13 clade with three present-day individuals, one Kenyan Bantu and two Dinka from Sudan (Figure 4). These relationships concur with those previously presented based on autosomal variants, with ancient South African individuals being more closely related to the San [27, 22], and Kenyan Pastoral Neolithic individuals having substantial Dinka-related ancestry (∼ 40%) with the remaining ancestry coming from North Africa and the Levant [24].

Regarding the B2-M82 lineages, in our dataset these are mostly composed of Biaka, Mbuti and San individuals. pathPhynder allowed further resolution of the lineage assignment of multiple ancient samples to B2b1-M192, including the second Shum Laka individual who was previously assigned to the B2b-M112 lineage (Figure 4). The reported autosomal affinity of Shum Laka samples with central African hunter-gatherer populations fits with this assignment to the B2 clade. We note, however, that in our dataset this sample’s Y chromosome is closest to that of a present-day Luhya individual from Kenya which carries a B2b1 lineage, rather than to those of the sampled Biaka, Mbuti and San.

Three samples from Malawi (two Malawi Fingira 6100BP and one Malawi Hora 9000BP), previously assigned to the BT-M91 macro-haplogroup [22], were in the present analysis further refined to B2b1-M192, B2b1b-M6557 and B2b1a1-M8349 (Supplementary Table S2). The connection between the Y-chromosome lineages of ancient and present-day South African hunter-gatherer populations corroborates the finding that San-related ancestry related was widely distributed in the past, and composed a large part of the ancestry of Malawi hunter-gatherers, previously reported based on autosomal data [22].

Sample Kenya Kakapel 3900BP which was previously assigned to the CT-M168 clade [28], was observed to be ancestral to this lineage, and was instead assigned to B2b1a1-M8349, shared with present-day Mbuti and Biaka samples (Figure 4). Fittingly, this sample in particular was observed to share substantial autosomal genetic ancestry with the Mbuti [28].

The next clade on which we will focus is E1b1b1a1-M78, which has a broad geographical range which encompasses North and East Africa, Europe and Western Asia [32]. All Morocco Iberomaurusian were positioned in this clade [25] as well as a Jordan Pre-pottery Neolithic B (PPNB) individual (Figure 5A). A single East African Pastoral Neolithic sample was placed in the E1b1b1a1a1b-V32 clade, together with a Luo, a Luhya and an Iranian individual. The Egypt Ptolemaic sample, a Late Stone Age and a Pastoral Neolithic individual were placed in the E1b1b1a1b2-L677 clade, which can be found in the present day in the Horn of Africa and Egypt [32]. In our data, this clade is represented by Palestinians, one Bedouin, two Balochi and one Iranian.

Three Moroccan Early Neolithic samples carry E1b1b1b1-M310.1 lineages [17], with the lineage of sample IAM.5 further refined to E1b1b1b1a1 ∼ -PF2535 with a single supporting marker. In our analysis, they were placed ancestrally to present-day Mozabite and Saharawi North African lineages (Figure 5B), which is in agreement with the finding from autosomal analyses that these samples comprised an endemic Maghrebi element still retained in present-day North African populations [17].

Apart from those described above, the majority (n=9) of other East African Pastoral Neolithic samples were placed in E1b1b1b2b-CTS10880 lineages (Figure 5C), a sub lineage of E1b1b1b2-Z830 found in the Levantine proto-agriculturalist Natufians and a pre-pottery Neolithic B Levantine sample [30], sister to E1b1b1b2a-Z1145 lineages still found in the Middle East. This increases the Y chromosome support for the model that pastoralism was introduced into East Africa with population movements from the Levant [33]. In our dataset, this clade is represented by Bantu from Kenya and South Africa and one Maasai. Additionally, one early pastoral and two Pastoral Iron Age individuals from Tanzania were placed in the E2a-M41 clade (Supplementary Figure S5).

## Discussion

We present the pathPhynder workflow which can efficiently assign informative variants to branches of phylogenetic trees and then use this variation for ancient DNA sample placement. We demonstrate the utility of our approach by placing aDNA samples into a reference Y-chromosome tree, in many cases leading to increased phylogenetic resolution. Our method works with routinely used formats in aDNA analysis (VCF and bam files) and does not require alignments in the fasta format, which for large data sets can be computationally expensive and time-consuming to generate. Furthermore, our method is ancient DNA-aware and can filter out potentially deaminated sites and other mismatches which are known to occur in ancient sequences and that can compromise the accuracy of phylogenetic inference. Indeed, we show that placement methods which rely on likelihoods are very negatively impacted by deamination, but that our filtering approach can effectively remove such errors and obtain correct placements (Supplementary Figure S4).

We provide two distinct methods for phylogenetic placement: a very fast maximum likelihood and the best path method, which is slightly slower. While the former is suitable for placing hundreds of samples in a matter of minutes, the latter can provide a highly detailed output containing information about the SNPs supporting or in conflict with query sample placement. This is particularly important for ancient DNA samples because they commonly diverge from the present-day tree at internal branches, in which case they will have a mixture of ancestral and derived genotypes at the SNPs defining this branch. Our tools allow the user to examine this pattern, which is not directly accessible using standard likelihood placement methods.

For Y-chromosome analysis in particular, the majority of aDNA studies rely on a catalogue of known haplogroup-defining SNPs maintained by ISOGG, an extremely useful resource which compiles and curates variation obtained from multiple studies. However, maintaining a SNP database requires extensive curation efforts which is time-consuming and laborious, and therefore, there is a lag between the generation of new data and incorporation of this novel variation into databases. Our method offers a quick solution for immediately making use of uncatalogued variation as new data sets emerge.

On the other hand, in some cases ISOGG SNPs provide additional resolution, because 42,863 ISOGG variants are not included in our dataset. There are multiple reasons for this: 1) because the 2,014 individuals included in our reference tree fail to capture all the lineages listed in the ISOGG database; 2) we restricted our data set to the ∼ 10.3 Mb regions of the Y-chromosome where variants can be called unambiguously, as recommended by Poznik et al. [4]. If the same filter is applied to ISOGG, this results in the exclusion of 7,694 SNPs; 3) even within this region certain other SNPs which are present in ISOGG may not have been genotyped in the reference VCF file; 4) a small subset of SNPs were not assigned to reference tree branches due to multiple mutations, genotyping error, or possibly inaccuracies in the tree topology, resulting in the presence of the derived allele at unrelated branches of the tree and consequently in low likelihood for SNP assignment and exclusion from analysis.

The incorporation of all ISOGG data in our analyses is not trivial. A subset of the variants listed by ISOGG have provisional status and may be repositioned to different locations of the Y-chromosome tree as additional data emerge. Additionally, as the tree topology grows in complexity and new branches emerge, SNPs previously defining the same haplogroups (i.e. positioned along the same branch), may be assigned to different branches, which also leads to modifications in haplogroup names. With pathPhynder, these issues do not occur because it uses a phylogeny which derives from the genotype data itself, making sample placement into the phylogenetic tree straightforward and independent of previous annotations, which may be incorrect and subject to change.

Our workflow can be applied to any haploid data set, including, but not limited to, the Y-chromosome and the mitochondria, and can also be used for phylogenetic placement of environmental DNA samples into pre-estimated trees. Future applications could include examining ancient Y-chromosome and mitochondrial lineages in ancient cattle [34], wolves and dogs [35, 36], for which large turnovers have occurred.

## Conclusion

The highly degraded nature and frequent low coverage of aDNA sequences creates difficulties for joint phylogenetic analyses of past and present-day populations. In the present work, we address this issue by presenting the pathPhynder workflow, which can efficiently identify and assign branch informative SNPs to the edges of a tree and use this variation for aDNA sample phylogenetic placement. Our method is aDNA-aware and scales well to the dimensions of current data sets, being able to process hundreds of samples in a few minutes. We apply our method to ancient Y-chromosome lineages and demonstrate that it frequently provides increased lineage resolution. pathPhynder will be useful for highly resolved lineage determination of low coverage ancient DNA data, opening new possibilities for the analysis not only of ancient human samples, but also of sequences belonging to other species, including those generated as part of metagenomic and environmental DNA experiments.

## Material and Methods

### Y-chromosomal data

Whole-genome sequenced present-day Y-chromosomal data from 1208 males from [15] was complemented by two Y-haplogroup A00 samples from [21], 41 from [20], 16 from [37] and 1071 samples from the low coverage 1000 Genomes Project dataset [14]. These were combined with 10 ancient samples from [20, 38, 29, 39, 40, 41, 42]. Genotype calling, filtering and Y haplogroup prediction are described in detail in [15]. Additionally, 334 samples from the 1000 Genomes Project were removed due to *≥* 10% missing data across the ∼ 10.3 Mb analysed Y-chromosomal regions. The vcf files of samples mapped to GRCh37 were lifted over to GRCh38 using picard (v2.7.2) (http://broadinstitute.github.io/picard/), followed by merging with the rest of GRCh38-based data using bcftools (v1.8) [43]. Modern samples from the [20] dataset were filtered for minimum read depth of 3, while no minimum depth filter was applied to the 1000 Genomes Project, Wong et al. 2017 and ancient samples due to lower coverage. Lastly, sites with 5% of missing calls across samples were removed. In the final dataset of 2,014 males a total of 9,832,836 sites remained, including 121,966 variant sites. The maximum likelihood Y-phylogeny including 2,014 samples and 121,966 variant sites was inferred using RAxML (v8.2.10) with the GTRGAMMA substitution model [44]. A complete list of the individuals in the reference dataset is available at Zenodo (DOI:10.5281/zenodo.4332182).

We downloaded previously catalogued ISOGG variants from https://isogg.org/tree/, as available on 03/08/2020, restricting our analysis to biallelic SNPs.

### Ancient DNA query sample placement into the Y-chromosome reference tree

In order to place ancient samples into the reference phylogenetic tree, we first assigned variants present in the reference VCF file using phynder. The resulting branch assignments were processed by pathPhynder using the ‘prepare’ step, which prepares bed format files for calling variants in the ancient samples, as well as producing an annotated sites file including information about the haplogroup defining variants (extracted from ISOGG 2019-2020 version), if any, and at which branch they occur.

We then ran pathPhynder’s ‘pileup and filter’ step to generate a pileup using samtools [43] at the informative sites identified with phynder and filtered these with default parameters, i.e. requiring at least that 70% of reads support a single genotype (-c 0.7), and filtered the resulting calls using the ‘default’ mode (-m), which removes potentially deaminated variants from analysis.

The next step is ‘choose Best Path’ in which the tree is traversed and query sample genotypes are evaluated in terms of support or conflict with every branch of the tree, the best path containing the highest number of support markers is chosen, as well as the best position in which to place the ancient sample in the tree. This step generates a plot indicating the best path and various tables which show detailed information about SNP and haplogroup status for each ancient sample.

The last step adds the ancient samples to tree and produces a newick file and a plot with the reference phylogeny which includes the query sample placements.

Results were plotted using the R programming language [45], and the R packages phytools [46], ape [47] and ggplot2 [48].

## Supplemental Data

Supplemental Data include 5 figures and 2 tables.

## Acknowledgments

R.M. was supported by an EMBO Long-Term Fellowship (No. ALTF 133-2017). P.H. was supported by Estonian Research Council Grant PUT1036. B.D.S. was supported by the Wellcome 4-year PhD programme in Mathematical Genomics and Medicine (code PFHZ/157). R.M. and R.D. were also supported by Wellcome grant WT207492. We thank David Poznik for advice related to the Y-chromosomal data belonging to the 1000 Genomes Project. We also thank Toomas Kivisild, Chris Tyler-Smith, Daniel Bradley and Lara Cassidy for early discussions related to this work.

## Data and Code Availability

pathPhynder can be downloaded from https://github.com/ruidlpm/pathPhynder/. We provide a VCF file of the ‘BigTree’ Y-chromosome dataset, aligned to GRCh37 and to GRCh38, and a reference phylogenetic tree for sample placement at Zenodo (DOI:10.5281/zenodo.4332182). All data analysed in this work is publicly available.

## Authors Contributions

Conceptualization, R.M. and R.D. ; Methodology, R.M., B.D.S. and R.D.; Software, R.M., B.D.S. and R.D., Formal Analysis, R.M., B.D.S., P.H. and R.D.; Investigation, R.M., B.D.S., P.H. and R.D.; Resources, R.M., B.D.S., P.H. and R.D.; Data Curation: R.M. and P.H.; Writing, Review & Editing, R.M., B.D.S., P.H. and R.D.; Visualization, R.M. and R.D.; Project administration, R.M. and R.D.; Funding Acquisition, R.M. and R.D.; Supervision, R.D.

---

**Table 1.**

| **Context** | **Number of samples** | **Study** |
| --- | --- | --- |
| Neolithic Morocco and Iberia | 7 | Fregel et al., 2018 [17] |
| Malawi, Tanzania and South Africa | 6 | Skoglund et al., 2017 [22] |
| Ptolemaic Egypt | 1 | Schuenemann et al. 2017 [23] |
| Later Stone Age to Iron Age Kenya and Tanzania | 27 | Prendergast et al., 2019 [24] |
| Pleistocene North Africa | 6 | van de Loosdrecht et al., 2018 [25] |
| Late Stone Age to Iron Age Cameroon | 2 | Lipson et al., 2020 [26] |
| Stone Age South Africa | 2 | Schlebusch et al., 2017 [27] |
| Late Stone Age to Iron Age Kenya, Congo and Botswana | 12 | Wang et al., 2020 [28] |
| Late Epipaleolithic to Early Bronze Age Jordan and Israel | 15 | Lazaridis et al., 2016 [30] |
| Ethiopia ~2500BC | 1 | Gallego-Llorente et al., 2015 [29] |

