## Supplementary Material for "Placing ancient DNA sequences into reference phylogenies"

### Supplementary Text 1 - Likelihood-based placement of query sequences into fixed phylogenetic trees

#### Overview

Starting with a fixed phylogenetic tree, or a 'reference tree', the main objective of the method is to place a new sample, which we call the 'query', into the tree without changing the original topology, using sequence data such as mapped ancient DNA reads. To do this, we use a likelihood approach to calculate the probability of the query attaching at each edge. We do not, however, compute the length of the new branch on which the query lies. This length would depend almost exclusively on the sites differing in the query from all those on the existing tree, which are relatively vulnerable to noise in the data. Since our algorithm is designed primarily for ancient DNA, which has higher levels of sequence noise, we do not attempt to ascertain private variants.

The input to the algorithm is a phylogenetic tree in the form of a .newick file, a VCF file containing genotype data for all the tips in the tree and a second VCF file, containing the genotypes of the query samples to be placed in the tree. A query sample VCF file can be prepared from BAM files using pathPhynder. The output is the best branch assignment and, for suboptimal branch assignments within a specified likelihood ratio of the optimum, their respective relative likelihood scores and posterior probabilities under a uniform branch prior. We also optionally report the lowest branch in the tree for which the posterior probability of being assigned to that branch or one below it is greater than $(1 - p)$, where $p$ is a user defined threshold.

#### Likelihood calculation

Ideally, we would like to be able to test the placement of the query at every possible location on each edge. However, since edges are continuous, this is an infinite space and even a discrete approximation on each edge is computationally costly in large trees. We avoid this complexity by an approximation equivalent to assuming that there are no mutations on the edge where the query attaches, so that the position of the query on the edge is irrelevant. For mutations that are indeed on this edge, this approximation corresponds to pushing them up to just after the parent node if the query contained the mutation, or pushing them down to just before the child node if the query did not contain the mutation. We find this approximation to have negligible effect on the results.

In order to calculate the likelihood $L$ of the tree with the query on a given edge $e$, we use Felsenstein's pruning algorithm [1]. This is the standard likelihood recursion on trees, also known as belief propagation or message passing in the field of graphical models.

We only consider biallelic sites, although as discussed below we do differentiate between transition sites (C↔T or A↔G), which have a higher mutation rate, and transversion sites. We allow for possible errors in the primary data by setting the likelihood of the data at the tip as $0.999$ for the recorded allele, and $0.001$ for the alternative allele. For missing data the likelihood at the tip is $1$.

For each edge $e$ with length $l_e$ we calculate the probability $p_{\text{flip}}(e)$ that an allele will change over the edge, and the probability $p_{\text{stick}}(e)$ that it will stay the same, as

$$p_{\text{flip}}(e) = (1 - \exp(-\mu l_e))/2$$
$$p_{\text{stick}}(e) = (1 + \exp(-\mu l_e))/2$$

for two mutation rates $\mu$, one for transitions and one for transversions, by default setting the transition rate to 4 times the transversion rate, which is the approximate ratio observed in humans, though this can also be set by the user as a parameter if desired.

Next we calculate for each edge two likelihoods per site. First, the likelihood $L_{\text{below}}(e, x)$ of the data below edge $e$ conditional on the allele at the bottom of the edge having value $x$, by applying the recursion up the tree from leaves to the root

$$L_{\text{below}}(e, x) = \left(L_{\text{below}}(e_L, x)p_{\text{stick}}(e_L) + L_{\text{below}}(e_L, \bar{x})p_{\text{flip}}(e_L)\right)\left(L_{\text{below}}(e_R, x)p_{\text{stick}}(e_R) + L_{\text{below}}(e_R, \bar{x})p_{\text{flip}}(e_R)\right)$$

where $e_L$ and $e_R$ are the left and right edges below $e$, and $\bar{x}$ is the alternative allele to $x$. Second, the likelihood $L_{\text{above}}(e, x)$ of all the other data at the site on leaves that are not below edge $e$, conditional on the allele at the top of the edge having value $x$, via the recursion down the tree from the root

$$L_{\text{above}}(e, x) = \left(L_{\text{above}}(e_P, x)p_{\text{stick}}(e_P) + L_{\text{above}}(e_P, \bar{x})p_{\text{flip}}(e_P)\right)\left(L_{\text{below}}(e_S, x)p_{\text{stick}}(e_S) + L_{\text{below}}(e_S, \bar{x})p_{\text{flip}}(e_S)\right)$$

where $e_P$ and $e_S$ are the parent and sibling edges of $e$, and $L_{\text{above}}(e_P) = 1$ for the edges immediately below the root. Assuming there are no mutations on the edge $e$ as mentioned above, we can then calculate the likelihood of both the top and bottom of the edge $e$ having value $x$ as

$$L(e, x) = L_{\text{above}}(e, x)L_{\text{below}}(e, x)$$

These calculations are all carried out in log space. We then add log likelihoods across sites to determine the log likelihood of the tree with the query placed on edge $e$. We store site patterns so that if two sites have the same pattern of data in their leaves, we only do the likelihood calculations once.

The method does not require the assignment of variants to edges, but sometimes these hard assignments are desired output. Variant assignment is therefore determined as follows. If there were indeed a mutation on the edge $e$ from $x$ to $\bar{x}$, we would calculate the likelihood $L(e, x)$ as

$$L(e, x) = L_{\text{above}}(e, x)L_{\text{below}}(e, \bar{x})$$

We can then choose the variant assignment based on maximum likelihood. A variant may be assigned to the root edges, in which case it will have equal likelihood of belonging on either one. To make a decision in these cases, we determine the maximum likelihood ancestral node state and then assign the variant to the appropriate branch.

Although we have described these algorithms for rooted trees, the code also calculates likelihoods correctly for unrooted trees which can be represented in .newick files via a “root” node with three descendants.

Optionally we can force the tree to be ultrametric, that is, to have edge lengths such that all the tips are the same distance to the root. This is done by balancing the tree, recursively scaling the left and right subtrees of a node so that they both have the same average height, setting this to the mean of their original average heights.

Table S1: List of published samples of unresolved haplogroup assignments reanalysed with pathPhynder.

| **Study** | **sampleID** | **population** | **reported haplogroup** | **newly assigned haplogroup** |
|---|---|---|---|---|
| Feldman et al 2019 [2] | ASH008.A0101 | Ashkelon Iron Age 2 | BT | J1a2a1a2d2b |
| Flegontov et al., 2019 [3] | I7342_d | Old_Bering_Sea | BT | Q1a1b |
| | I0721 | Paleo-Aleut | CT | P1orK2b2a |
| Lazaridis et al., 2016 [4] | I1685 | Israel_Raqefet_M_Natufian | CT | E1b1b1 |
| | I1700 | Jordan_PPNB | CT | H2 |
| Mathieson et al., 2017 [5] | I2407 | Globular_Amphora_Poland | BT | F |
| | I2426 | Balkans_Chalcolithic | CT | F |
| | I2435 | Globular_Amphora_Poland | CT | I2a |
| Narasimhan et al., 2018 [6] | I12257 | Ksirov_H_Kushan | n/a | R |
| | I12293 | Ksirov_H_Kushan | n/a | G2a2b1 |
| | I12980 | Loebanr_IA | n/a | F |
| | I1945 | Ganj_Dareh_N | n/a | R2a |
| | I1949 | Ganj_Dareh_N | n/a | R2a |
| | I3389 | Central_Steppe_MLBA | n/a | R1a1a1b2 |
| | I6117 | Gonur1_BA | n/a | Q1a1a1b2 |
| | I11562 | Parwak_Medieval | BT | BT |
| | I4319 | Kairan_MLBA | BT | P1orK2b2a |
| | I5762 | Kairan_MLBA_LowCov | BT | R1a1a1 |
| | I6120 | Gonur1_BA | BT | F |
| | I7101 | Gonur1_BA | BT | R2a2 |
| | I11042 | Gonur1_BA | CT | J |
| | I12140 | Katelai_IA | CT | CT |
| | I2328 | Hajji_Firuz_C | CT | J |
| | I4158 | Bustan_BA | CT | F |
| | I6119 | Gonur1_BA | CT | Q1b |
| | I6312 | Gonur1_BA | CT | R1b1a1b1a1 |
| | I7170 | Gonur1_BA | CT | CT |
| | I8527 | Geoksyur_EN | CT | J1a |
| | I11458 | Shahr_I_Sokhta_BA2 | F | P1orK2b2a |
| Ning et al., 2019 [7] | M4 | Shirenzi | n/a | R1a1 |
| | M8R1 | Shirenzi | n/a | O2a2 |
| Skoglund et al., 2017 [8] | I2966 | Malawi_Hora_9000BP | BT | B2b1b |
| | I4427 | Malawi_Fingira_6100BP | BT | B2b1a1 |
| | I4468 | Malawi_Fingira_6100BP | BT | B2b1 |

Table S2: List of African and Middle Eastern ancient DNA samples placed into the reference Y-chromosome tree. The tilde indicates lineages with uncertain position in the ISOGG Y-chromosome tree.

| Study | sampleID | population | reported haplogroup | newly assigned haplogroup |
|---|---|---|---|---|
| | I1414 | Jordan_PPNB | E | E1b1b1b2 |
| | I1072 | Israel_Raqefet_M_Natufian | E1b1b1b2 | E1b1b1b2 |
| | I1707 | Jordan_PPNB | T | T1a |
| | I1730 | Jordan_EBA | J | J2b1 |
| | I1705 | Jordan_EBA | J1 | J1a2a1a2d2b2b∼ |
| | I0861 | Israel_Raqefet_M_Natufian | E1b1b1b2 | E1b1b1b2 |
| | I1415 | Jordan_PPNB | E1b1b1 | E1b1b1 |
| Lazaridis et al., 2016 [4] | I1710 | Jordan_PPNB | E1b1b1 | E1b1b1a1 |
| | I1727 | Jordan_PPNB | CT | CT |
| | I1069 | Israel_Raqefet_M_Natufian | E1b1 | E1b1 |
| | I1700 | Jordan_PPNB | CT | H2∼ |
| | I0867 | Israel_PPNB | H2 | H2∼ |
| | I1690 | Israel_Raqefet_M_Natufian | CT | CT |
| | I1416 | Levant_PPNB | CT | CT |
| | I1685 | Israel_Raqefet_M_Natufian | CT | E1b1b1∼ |
| | TAF014 | Morocco_Iberomaurusian | E1b1b1a1 | E1b1b1a1 |
| | TAF013 | Morocco_Iberomaurusian | E1b1b1a1 | E1b1b1a1 |
| van de Loosdrecht et al., 2018 [9] | TAF015 | Morocco_Iberomaurusian | E1b1b1 | E1b1b1a1 |
| | TAF010 | Morocco_Iberomaurusian | E1b1b1a1 | E1b1b1a1 |
| | TAF009 | Morocco_Iberomaurusian | E1b1b1a1 | E1b1b1a1 |
| | TAF011 | Morocco_Iberomaurusian | E1b1b1a1 | E1b1b1a1 |
| | I0595 | Kenya_400BP | E1b1b1b2 | E1b1b1b2b2∼ |
| | I9028 | South_Africa_2100BP | A1b1b2a | A1b1b2a |
| Skoglund et al., 2017 [8] | I2966 | Malawi_Hora_9000BP | BT | B2b1b∼ |
| | I4427 | Malawi_Fingira_6100BP | BT | B2b1a1∼ |
| | I9133 | South_Africa_2000BP | A1b1b2a | A1b1b2a |
| | I4468 | Malawi_Fingira_6100BP | BT | B2b1∼ |
| | IAM.5 | Morocco_EN | E1b1b1b1a | E1b1b1b1a1∼ |
| | IAM.4 | Morocco_EN_dup.IAM5 | E1b1b1b1 | E1b1b1b1 |
| | IAM.7 | Morocco_EN | n/a | E1b1b1b1 |
| Fregel et al., 2018 [10] | TOR.12 | Iberia_EN | n/a | BT |
| | KEB.6 | Morocco_LN | T1a | T1a1 |
| | KEB.7 | Morocco_LN | n/a | CT |
| | TOR.5 | Iberia_EN | G2a2b2a3a | G2a |
| | I12379 | EmurulaOlePolosCairns(GvJh122)_PIA/recent | E1b1b1b2b2a1 | E1b1b1b2b2a1a∼ |
| | I12384 | OlKalou_PN(other) | E1b1b1b2b2a1 | E1b1b1b2b2a1 |
| | I12391 | Kasiole2burial(GvJh54)_PIA | E1b1b1b2b | E1b1b1b2b2a∼ |
| | I12398 | RigoCave(GrJh3)_PN(Elmenteitan) | E1b1b1b2b2a1 | E1b1b1b2b2a1 |
| | I12533 | Prettejohn'sGully(GsJi11)_Earlypastoral | E2(xE2b) | E2a∼ |
| | I13762 | GishimangedaCave_PN(other) | E1b1b1b2b2a1 | E1b1b1b2b2a1 |
| | I13972 | GishimangedaCave_PN(other) | E1b1b1b2b2 | E1b1b1b2b2 |
| | I13977 | GishimangedaCave_PN(other) | E1b1b1b2b2 | E1b1b1b2b2 |
| | I13980 | GishimangedaCave_PN(other) | E1b1b1a1b2 | E1b1b1a1b2 |
| | I13983 | GishimangedaCave_PN(other) | BT(xCT) | B2b1 |
| | I8759 | NaishiRockshelter_PN(other) | E1b1b1b2b | E1b1b1b2b |
| | I8802 | DeloraineFarm(GqJh6)_IA | E1b1a1a1a1a | E1b1a1a1a1a3∼ |
| | I8808 | JawuoyoRockshelter_LSA | E1b1b1a1b2 | E1b1b1a1b2 |
| Prendergast et al., 2019 [11] | I8830 | NaivashaBurialSite_PN(other) | xBT | A1b1b2b |
| | I8892 | IlkekMounds/IlkekII(Gilgil)(GsJj66)_PIA | E2(xE2b) | E2a |
| | I8901 | KisimaFarm,CairnKFR-C4_PIA | E2(xE2b) | E2 - but E2a |
| | I8904 | KokurmatakoreCairn2(GdJn2)_PIA | E1b1b1(xE1b1b1b2b) | E1b1b1∼ |
| | I8923 | RigoCave(GrJh3)_PN(Elmenteitan) | E1b1b1b2b2 | E1b1b1b2b2 |
| | I8930 | WhiteRockPoint(GrJb2)_LSA | BT(xCT) | B2b1a1∼ |
| | I8758 | NaishiRockshelter_PN(other) | A1b(xA1b1b2a) | A1b1b2b2∼ |
| | I8804 | KeringetCave(GrJg4)1_PN(other) | A1b1b2 | A1b1b2b∼ |
| | I8919 | NaivashaBurialSite_PN(other) | A1b1b2b | A1b1b2b2∼ |
| | I8874 | Cole'sBurial(GrJj5a)_PN(other) | E1b1b1a1a1b1 | E1b1b1a1a1b1a∼ |
| | I8809 | KisimaFarmA5(PorcupineCave)_PN(other) | E1b1b1b2b2a1 | E1b1b1b2b2a1a∼ |
| | I8918 | NaivashaBurialSite_PN(other) | E1b1b1b2b2a1 | E1b1b1b2b2a1a∼ |
| | I8920 | NaivashaBurialSite_PN(other) | E1b1b1b2b2a1 | E1b1b1b2b2a1a1b∼ |
| | I8922 | RigoCave(GrJh3)_PN(Elmenteitan) | E1b1b1b2b2a1 | E1b1b1b2b2a1a1b∼ |
| Lipson et al., 2020 [12] | 4/A - I10873 | Shum Laka | B2b | B2b1∼ |
| | 2/SE II - I10871 | Shum Laka | A00 | A00 |
| Schlebusch et al., 2017 [13] | BAA | Ballito Bay A | A1b1b2 | A1b1b2b2∼ |
| | BAB | Ballito Bay B | A1b1b2 | A1b1b2b2∼ |
| | NYA003 | Kenya_Nyarindi_3500BP | E | CT |
| | LUK001 | Kenya_LukenyaHill_3500BP | E1b1b1b2b | E1b1b1b2b2a∼ |
| | HYR002 | Kenya_HyraxHill_2300BP | E1b1b1b2b | E1b1b1b2b2a1 |
| | MOL001 | Kenya_MoloCave_1500BP | E1b1b1b2b | E1b1b1b2b2a1 |
| | KPL001 | Kenya_Kakapel_3900BP | CT | B2b1a1∼ |
| Wang et al., 2020 [14] | KIN002 | Congo_Kindoki_230BP | E1b1a1a1d1a2 | E1b1a1a1a2a1a3b1a2 |
| | KIN003 | Congo_Kindoki_150BP | E | CT |
| | KIN004 | Congo_Kindoki_230BP | R1b1 | R1b1b2a |
| | NGO001 | Congo_NgongoMbata_220BP | n/a | E1b1a1a1a1c1a |
| | TAU001 | Botswana_Taukome_1100BP | E1b1a1 | E1b1a1∼ |
| | XAR001 | Botswana_Xaro_1400BP | E1b1a1a1c1a | E1b1a1a1a1c1a1a3 |
| | XAR002 | Botswana_Xaro_1400BP | E1b1b1b2b | E1b1b1b2b2 |
| Schuenemann et al. 2017 [15] | JK2888 | Egypt_Ptolemaic | E1b1b1a1b2 | E1b1b1a1b2 |

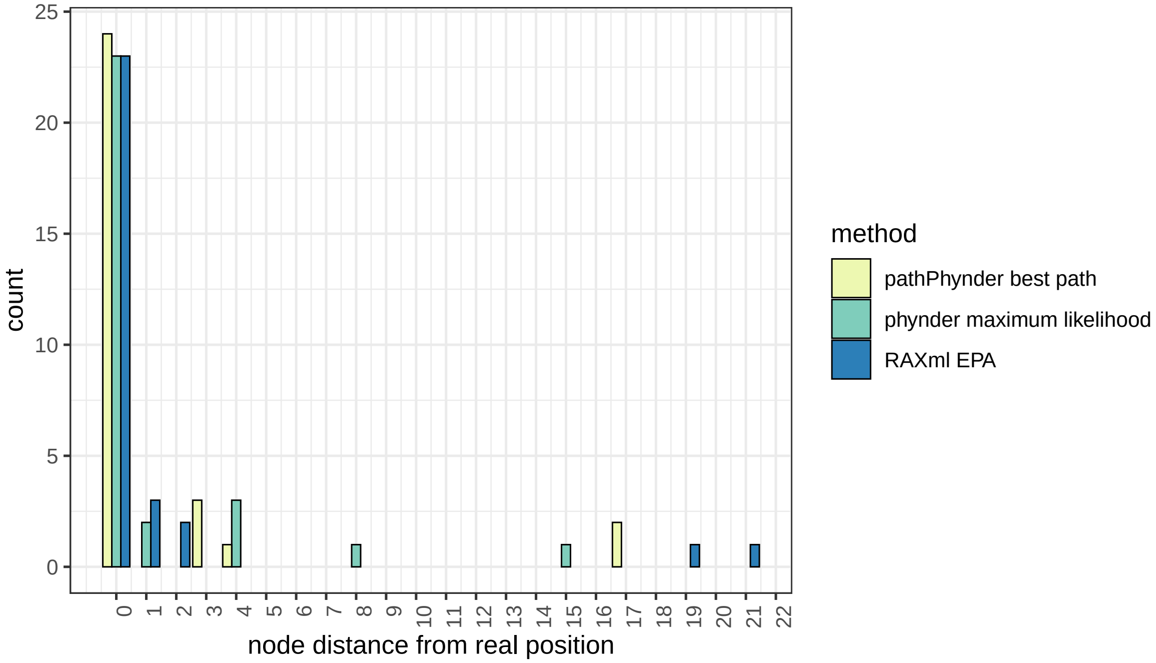


Figure S1: Comparison between three different methods for sequence placement into a reference phylogenetic tree. The x-axis indicates placement distance from the real position measured by the number of nodes which separate them.

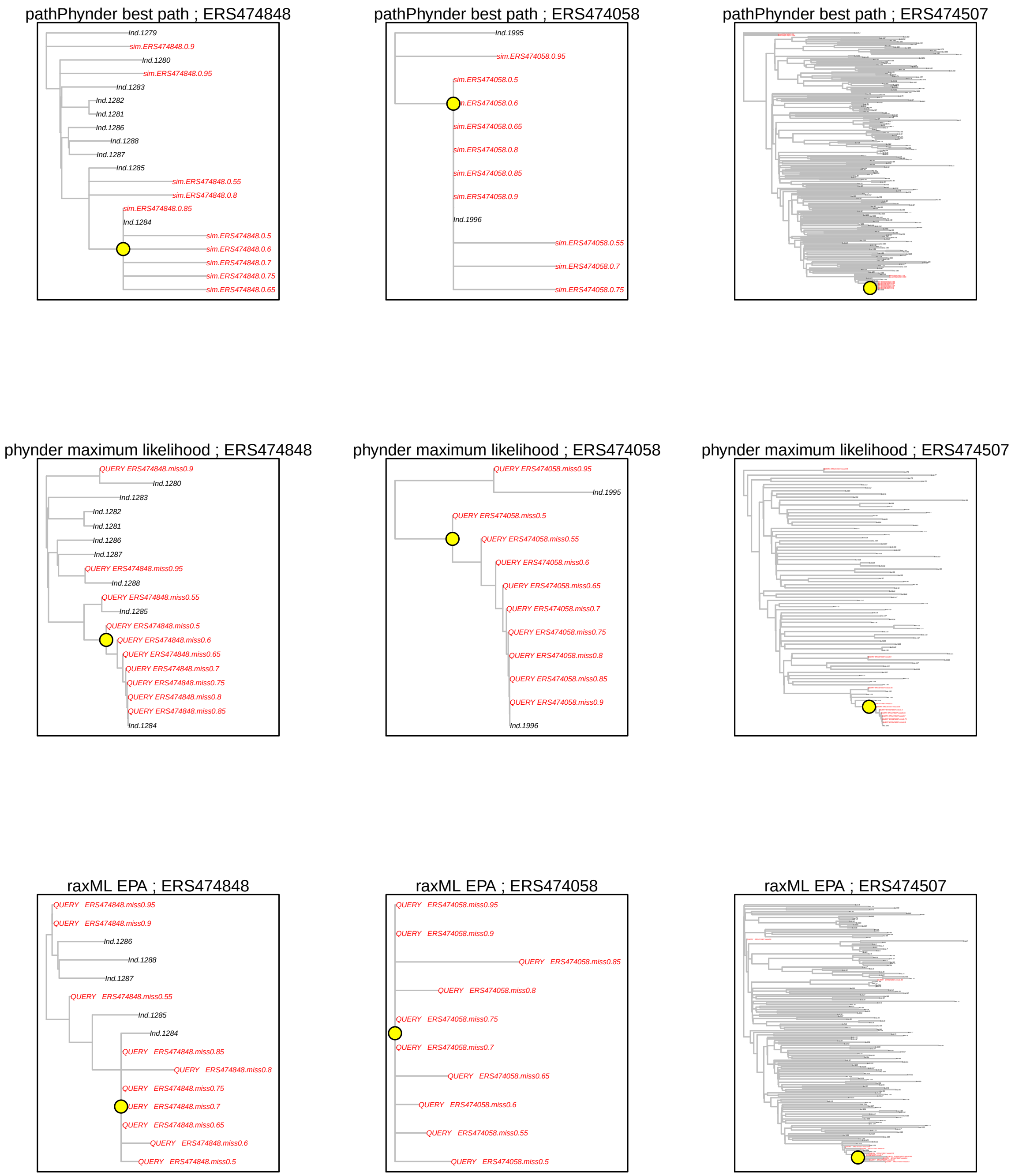


Figure S2: Comparison of the effect of missing data in query placement with RAxML EPA, phynder Maximum Likelihood and pathPhynder Best Path. The query samples are three individuals (ERS474848 - Bedouin, ERS474058 - Biaka, ERS474507 - Brahui) with artificially inserted missing genotypes at a proportion ranging from 0.5 to 0.95, for a total of 30 queries (coloured in red). The yellow circle indicates the correct location in the phylogenetic trees for sample placement.

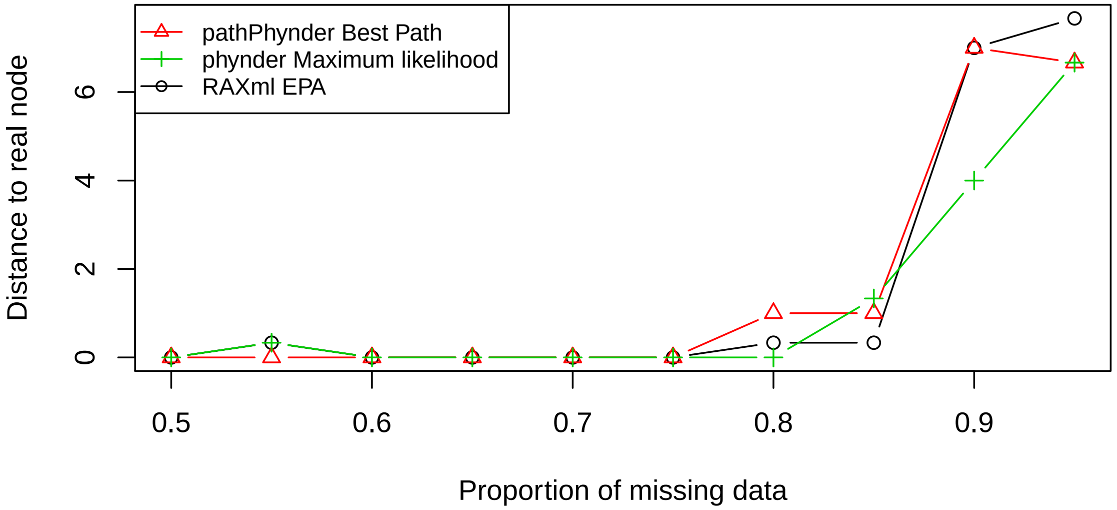


Figure S3: Comparison between three different methods for sequence placement into a reference phylogenetic tree. The x-axis indicates the proportion of missing data and the y-axis shows the mean distance from the real position across the three samples for each downsampling.

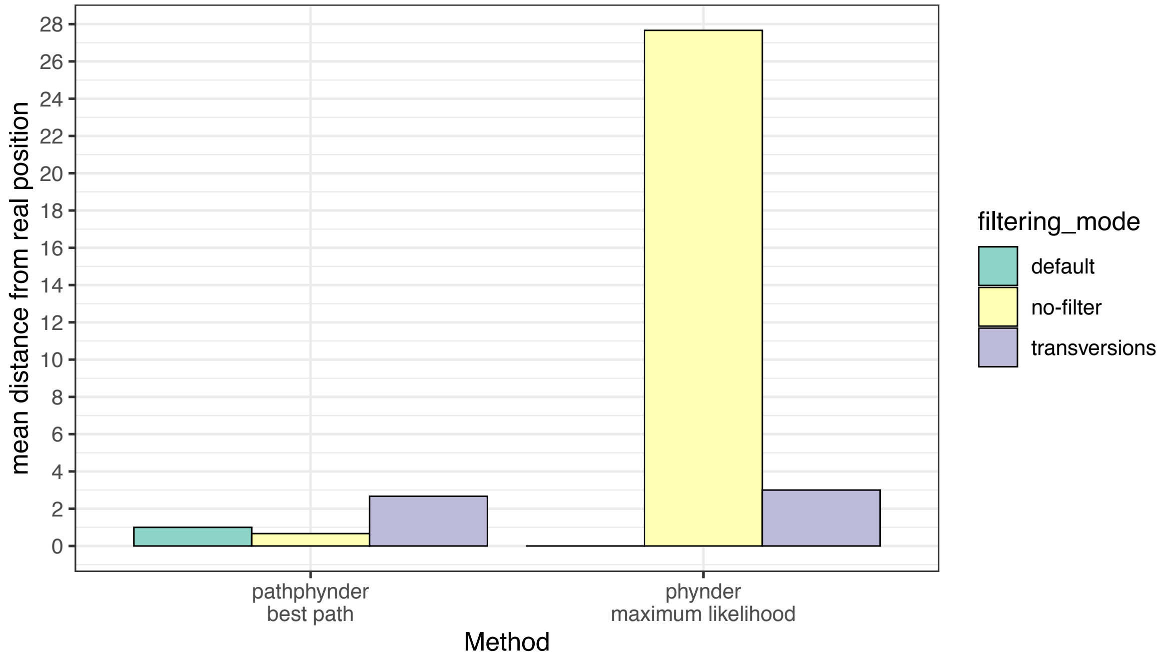


Figure S4: Comparison between the placement of the best path and maximum likelihood methods using different modes for filtering. In the default (conservative) mode, sites with potential deamination are removed, in the no-filter mode, they are kept, and the transversions mode excludes transitions from analysis.

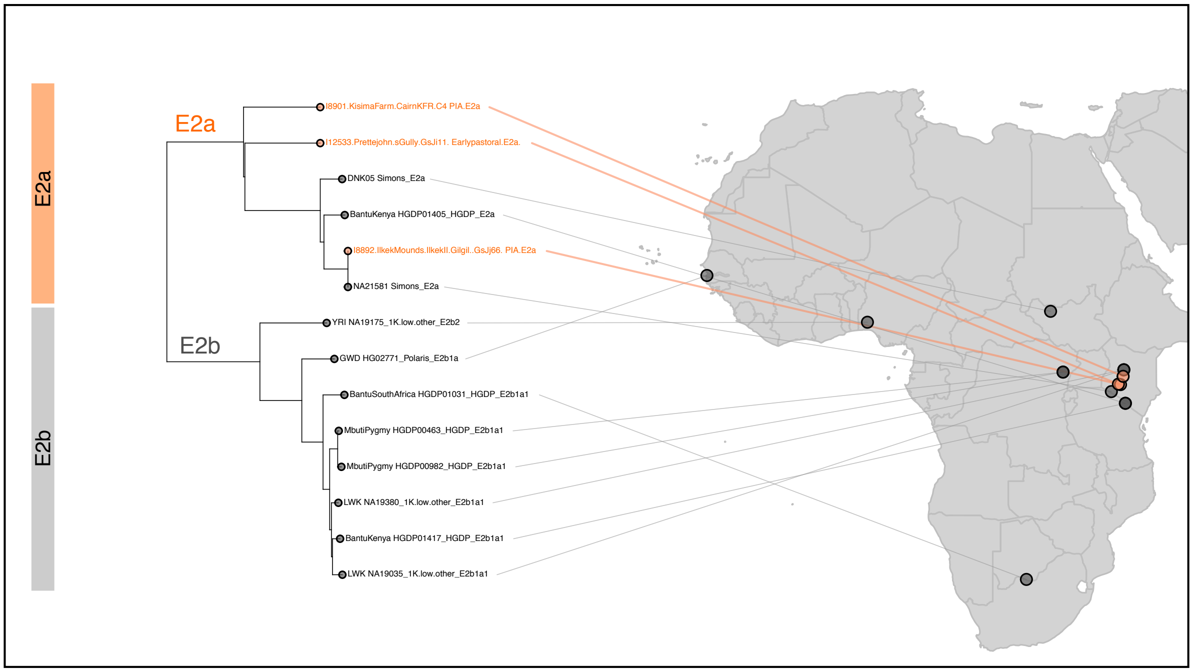


Figure S5: Placement of Early Pastoralist and Pastoralist Iron Age African samples in the E2b clade of the Y-chromosome tree.
